# Synchrony of Non-isochronous Signals in an Acoustically Communicating Katydid

**DOI:** 10.1101/2020.12.10.419101

**Authors:** Vivek Nityananda, Rohini Balakrishnan

## Abstract

The ability to entrain to auditory stimuli has been a powerful method to investigate the comparative rhythm abilities of different animals. While synchrony to regular (isochronous) rhythms is well documented, synchrony to non-isochronous stimuli, with multiple components at unequal time intervals, is rarer. Several katydid species with isochronous calls have been shown to achieve synchrony as part of their natural calling interactions in multi-individual choruses. Yet no study so far has demonstrated synchrony in any insect with a non-isochronous call. Using natural calling behaviour and playback experiments, we investigated acoustic synchrony and the mechanisms underlying it in the katydid species *Mecopoda* ‘Two Part Caller’. This species has a complex non-isochronous call consisting of a long trill followed by two or more chirps. We found that individual males synchronized trills and, to a lesser extent, chirps. Further investigation of trill synchrony showed that timing of trills is modified by external trills but not chirps. Chirp synchrony is modified by external chirps but also by trills. We suggest a two-oscillator model underlying synchrony in this species and discuss the implications for the evolution of acoustic synchrony.

## Introduction

Several animals have a striking ability to synchronize motor activity in response to external stimuli. The study of the comparative abilities of animals to do so has been an important field of research (Cook et al., 2013; Hasegawa et al., 2011; Jao Keehn et al., 2019; Patel et al., 2009; Schachner et al., 2009; Wilson and Cook, 2016). Experiments investigating these abilities often involve playback of artificial rhythmic stimuli to animals and studying their response as evidence of their rhythmic abilities. Rhythmic responses have, however also been demonstrated in studies of natural behaviour in other animals, including fireflies, frogs and katydids (bushcrickets) (Greenfield, 2002). Males of several katydid species produce calls to attract mates over a long distance (Alexander, 1967). In many of these species, males call together in group assemblages called choruses (Greenfield, 1994) and interact with each other acoustically to produce striking temporal patterns such as synchrony and alternation (Greenfield and Roizen, 1993a; Greenfield and Schul, 2008; Greenfield and Snedden, 2003; Hartbauer et al., 2005; Nityananda and Balakrishnan, 2007). Their natural behaviour thus serves as an ideal method by which to investigate their rhythmic abilities. Studies of acoustic synchrony in katydids have therefore typically focussed on the responses of individuals to natural stimuli both in the context of playback and in multi-signaller choruses.

Acoustic synchrony in katydids is often imperfect and the calls of individual males lead or follow calls of other males by a few milliseconds (Greenfield and Roizen, 1993b; Hartbauer et al., 2005; Nityananda and Balakrishnan, 2007). The mechanisms underlying this synchrony differ in different species and various models have been suggested to describe the mechanisms underlying both acoustic synchrony in katydids and the closely related phenomenon of synchronous flashing in fireflies (Buck et al., 1981a; Buck et al., 1981b; Greenfield, 1994). In species of the katydid genus *Mecopoda*, these mechanisms have typically been investigated using Phase Response Curves (PRCs), which represent the response of an internal oscillator when disturbed by single chirps at different phases of the calling cycle (Hartbauer et al., 2005; Nityananda and Balakrishnan, 2007; Sismondo, 1990). A typical PRC thus demonstrates how the animal increases or decreases the length of its calling period in response to external chirps. In the species *Mecopoda elongata,* the slope of the PRC seems to determine whether synchrony or alternation occurs (Sismondo, 1990). Simulations of duets based on PRCs also show that the male with the faster intrinsic chirp rate produces chirps that predominantly lead those of a male with a slower rate (Hartbauer et al., 2005). In the species *Mecopoda* ‘Chirper’, a similar approach showed that synchrony is enabled in this species through a combination of chirp-by-chirp resetting and a change in intrinsic rate (Nityananda and Balakrishnan, 2007).

Different mechanisms thus enable synchrony in different species. All species that have been studied, however, have had simple calls consisting of a single chirp repeated at a regular rhythm, that is, isochronous calls. Studies investigating rhythm perception in other animals have also focussed on their ability to perceive isochronous rhythms (Celma-Miralles and Toro, 2018). Evidence for entrainment to more complex rhythms has been shown for vocal learners like parrots (Patel et al., 2009; Schachner et al., 2009), but also more recently from sea lions (Cook et al., 2013). In katydids, synchrony has thus far not been reported from species that have more complex calls and the mechanisms that might underlie such synchrony remain unknown.

We here demonstrate acoustic synchrony between individuals of *Mecopoda* ‘Two Part Caller’, a katydid species with a complex non-isochronous call consisting of a trill, followed by two or more short chirps (Fig. 1). This is one of five southern Indian species belonging to the genus *Mecopoda* that look identical but have dramatically different songs (Nityananda and Balakrishnan, 2006). This song type is found in the evergreen forests of the state of Karnataka in South India and has a peak breeding season between the months of March and June. The call of this species consists of a verse, comprising a trill followed by a set of chirps (Fig. 1). Using acoustic playback experiments, we investigated acoustic synchrony and the possible mechanisms underlying it in this species.

**Fig. 1.**
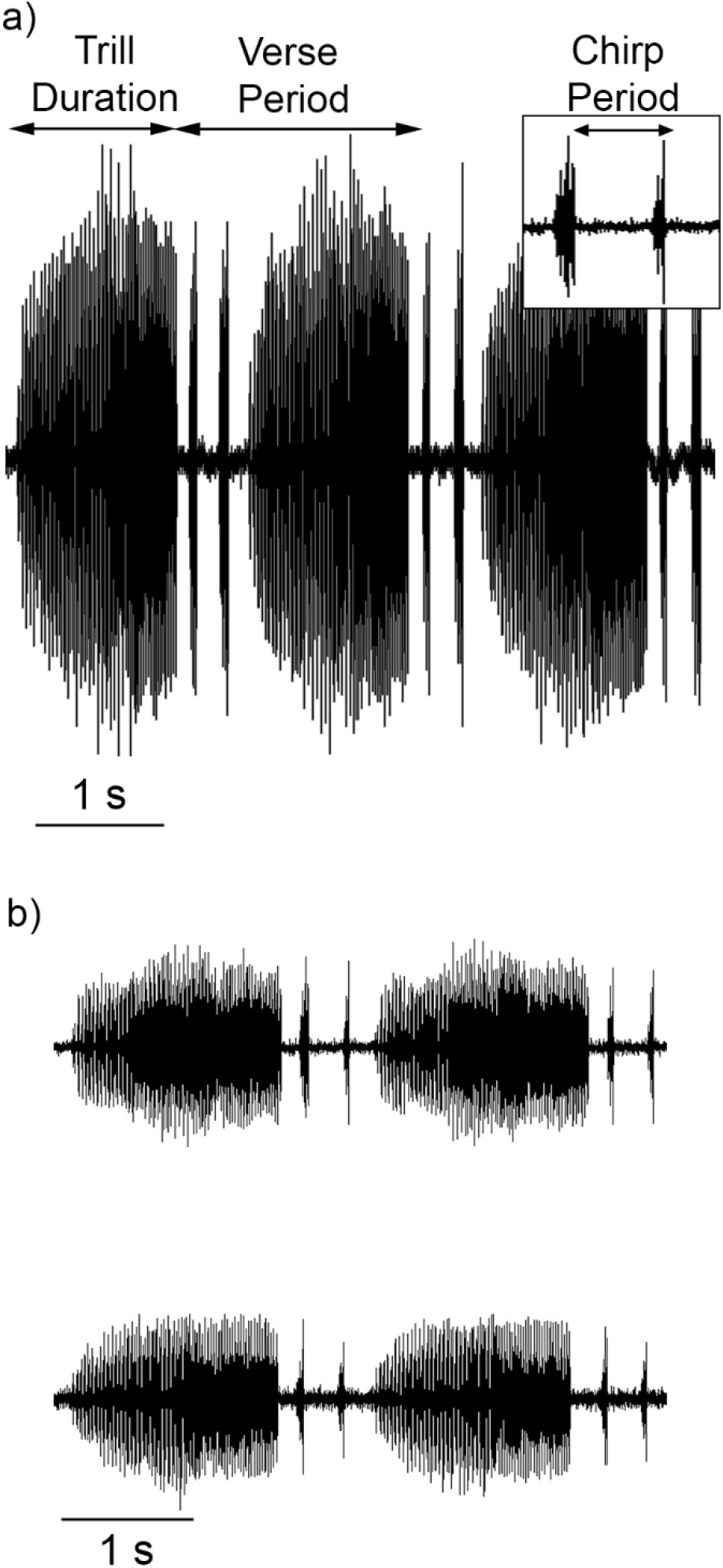
a) Oscillogram of the call of *Mecopoda* ‘Two Part Caller’ b) Oscillogram of two simultaneously calling males with synchronized trills. Inset shows a magnified view of the chirps,

## Materials and Methods

In all experiments, individual males were placed in acoustically transparent nylon mesh cages and the calling of the males and any stimuli played back were recorded on separate channels in an anechoic chamber (2 × 2 × 3 m) using tiepin microphones and custom-built amplifiers placed directly in front of the cages. The output of the microphones was digitized at a sampling rate of 16 kHz using an NI-DAQ AT-MIO-16E-2 card and the software Labview 6.0 (National Instruments Corporation, USA).

All playback experiments used stimuli taken from a previous recording of an individual ‘Two Part Caller’ male made using a Bruel and Kjaer Sound Level Meter (Type 2231with a ¼” microphone (4939: flat frequency response from 4 Hz to 70 kHz) and digitized at a sampling rate of 200 kHz using a NI-DAQ AT-MIO-16E-2 card and the software Labview 6. 0 (National Instruments Corporation, USA). Stimuli were played out at a sampling rate of 200 kHz using an NI-DAQ AT-MIO-16E-2 card and an Avisoft Ultrasonic Scanspeak speaker (frequency range 1-120 kHz) from a distance of 2 m. The stimuli were played out at 91 dB SPL (peak) 30 cm from the speaker measured using a Bruel and Kjaer Sound Level Meter type 2231 with a ¼ in 4939 microphone (frequency response 4-70 kHz).

### Song recording and analysis

All recordings were made at night, during the natural calling time of the animals. Solo recordings were made with individuals isolated in the anechoic chamber and duets were recorded with the two cages placed 2 m apart from each other in the anechoic chamber. During duets and the Phase Response Curve experiments, outputs were simultaneously obtained on two separate channels from the two microphones. Custom MATLAB (Mathworks, USA) programs were subsequently used to first obtain the time of call onsets and offsets and then to calculate the period, duration and phase relationships of the call elements. A Testo 110 thermometer (Testo Ltd., UK) was used to measure the ambient temperature during recordings. The mean temperature across all recordings was 22.23 °C (± 0.30 S. D.).

Phase relationships between trills were calculated according to the formula

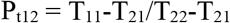

where T_11_ is the time of offset of the trill, T_21_ is the time of offset of the preceding trill of the partner and T_22_ is the time of offset of the following trill of the partner.

Phase relationships of the chirps were calculated with the same formula using chirp offsets instead of trill offsets. Phase values thus obtained were multiplied by 360° to obtain the phase angles. The x and y components of each of these angles were determined assuming each angle was a vector of unit length. The components for each duet were then summed together to obtain the x and y components of the mean vector for each duet. The angle and length of the mean vectors were then calculated (Batschelet, 1981). A V-test (Batschelet, 1981) was performed to test if the vectors were clustered around an angle of 0° which would indicate perfect synchrony. The mean trill and chirp periods during solos and duets were compared using unpaired t-tests at a significance level α = 0.05.

### Phase Response Curves

To obtain the Phase Response Curves (PRCs), a previously recorded stimulus was played out after the male initiated calling. The stimulus was played out at a random phase of the male’s calling, approximately once every 7 verse periods.

We obtained PRCs in response to playback of both trills (Trill PRCs) and chirps (Chirp PRCs). In both trill and chirp playback experiments, we looked at the effect on both trills and chirps and obtained separate PRCS for each with calculations as described below. We thus had four types of PRCs demonstrating the effect of trill playback on trills, trill playback on chirps, chirp playback on trills and chirp playback on chirps. For all four PRCs, the stimuli played out and the calls of the male were simultaneously recorded as described above. The times of offsets and onsets were obtained and phases were calculated as detailed below.

### The effect of trill playback on trills

For this PRC, the stimulus phase was determined according to the formula

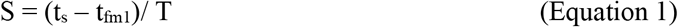

where S is the stimulus phase, t_s_ is the time of offset of the stimulus trill, t_fm1_ was the time of offset of the preceding trill of the male and T was the undisturbed verse period of the calling male calculated as the mean of three verse periods of the male before the playback (Fig. 2a). Stimulus phases close to 0 represent stimuli played out soon after the end of preceding trill and phases close to 1 represent stimuli played out close to the next trill.

**Fig. 2.**
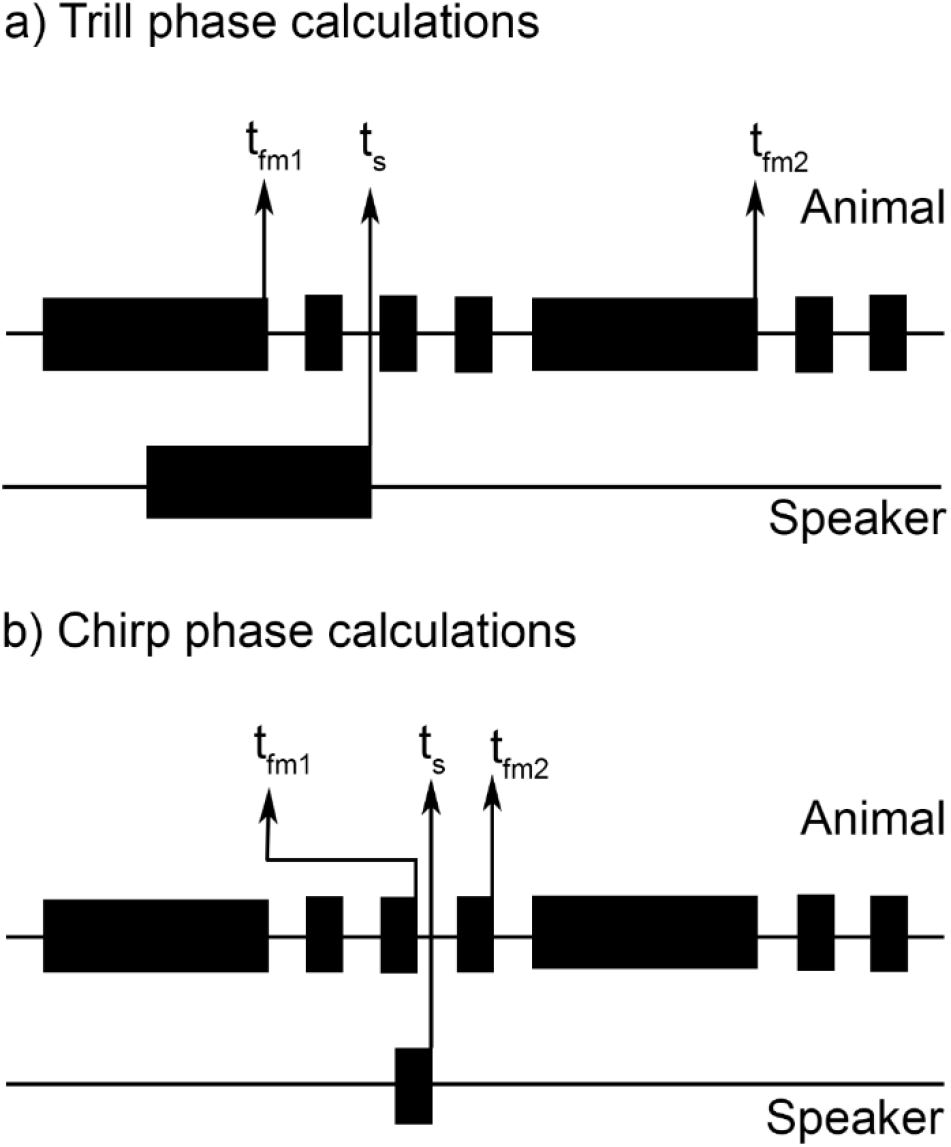
Phase calculations for the Phase Response Curves. Long black rectangles represent trills and shorter rectangles represent chirps. a) Trill phase calculations computed phase with respect to the offsets of the stimulus played back from the speaker and the *trills* of the calling animal. Note that the speaker stimulus here is depicted as a trill but in other experiments this was a chirp. b) Chirp phase calculations computed phase with respect to the offsets of the stimulus played back from the speaker and the *chirps* of the calling animal. Note that the speaker stimulus here is depicted as a chirp but in other experiments this was a trill. See the main text for calculations and further details.

The response phase was determined according to the formula

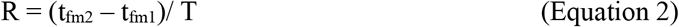

where R is the response phase.

t_fm1_ was the time of offset of the trill preceding the stimulus, t_fm2_ was the time of offset of the following trill of the male and T is the undisturbed verse period of the calling male calculated as the mean of three verse periods of the male before the playback (Fig 2a). Response phases close to 1 represent no change in period in response to playback and phases greater than 1 represent increases in verse period in response to playback.

### The effect of trill playback on chirps

For this PRC, the calculation were made as in Equations 1 and 2 except that t_fm1_ was the time of offset of the preceding chirp of the male, t_fm2_ was the time of offset of the following chirp of the male and T was the undisturbed chirp period of the calling male calculated as the mean of three chirp periods of the male before the playback. Note that when the period between two chirps included a trill, this chirp period was not used to calculate the undisturbed chirp period.

### The effect of chirp playback on trills

For this PRC, calculations were made using Equations 1 and 2 except that t_s_ was the time of offset of the stimulus chirp.

### The effect of chirp playback on chirps

For this PRC, the calculations were made as for the effect of trill playback on chirps except that t_s_ was the time of offset of the stimulus chirp (Fig 2b).

In all four cases, the Phase Response Curve (PRC) was obtained by plotting the response phase against the stimulus phase. The PRC gives us a picture of how the animal adjusts the timing of its next trill or chirp in response to stimulus playback at different phases in its calling cycle.

The means of three trill periods before and after the period disturbed by stimulus playback were calculated and compared using student’s t-tests with the significance level was Bonferroni corrected for multiple comparisons (Cabin and Mitchell, 2010). The tests were used to determine if there was a significant difference in trill period caused by playback of the stimulus. A similar comparison of means was made between the means of three trill periods before one stimulus playback and the next one approximately seven periods later. This enabled us to examine if the effects of stimulus playback on trill period were long-lasting.

### Response to chirpless calls

To investigate the influence of chirps on the synchronization of trills between males, we conducted a series of tests in which the response of males to calls without chirps was recorded. Two categories of experiments were conducted for each male: a positive control duet and a duet with chirpless calls. During the positive control duets, the recording played out to the male consisted of a standard ‘Two Part Caller’ verse of duration 1.89 (±.11 S.D.) s and a period of 2.05 (±.09) s followed by two chirps. During the chirpless duets, the same call was used with the chirps removed and replaced by silence. The call onsets and offsets were obtained and analyzed as previously described.

## Results

### Solos and duets

During solo calling, individuals (N = 22) produced verses with a mean period of 1.92 s (±.10 S.E., N= 236.77 ± 49.79 S.E verses per individual). These consisted of trills with a mean duration of 1.31 s (±.07 S.E.) followed by chirps of mean duration of 67.8 ms (± 0.36 S.E., N= 272.09 ± 82.99 S.E chirps per individual) (Figure 1a). The mean number of chirps following the trills in the solo bouts was 2.02 (± 0.065 S.E.).

In acoustic duets in the laboratory, individual males (N=18 duets, 36 males) synchronized their trills (N = 144.39 ± 26.24 S.E. trills per focal individual), with a mean vector angle of 7.41°, where an angle of 0° or 360° would represent perfect synchrony (Fig. 3a,c). The vector angles across all duets were within 56° of perfect synchrony (0°/360°) (Fig 3c). The length of the average mean vector across all duets was 0.92 (±.01 S.E.) and ranged from 0.76 to 0.99 indicating a high degree of synchrony (a vector length of 1 represents perfect synchrony). The distribution of mean vector angles was significantly different from a uniform distribution and the V-test confirmed that the angles were centred at 0° (V-test, v = 16.39, P = 2.35 * 10^-8^). Males also synchronized their chirps (N = 179.17 ± 42.32 S.E. chirps per individual); the distribution of chirp phase angles was also significantly different from a uniform distribution and centred around 0° (V-test, v= 11.52 P = 6.15*10^-5^, Fig. 3b, d). The degree and precision of synchrony, however, was less than that of the trills (Fig. 3d) with an average mean vector length of 0.64 (±.07 S.E.) and spread over a larger range of angles with a maximum angle of 121.99° from perfect synchrony.

**Fig. 3.**
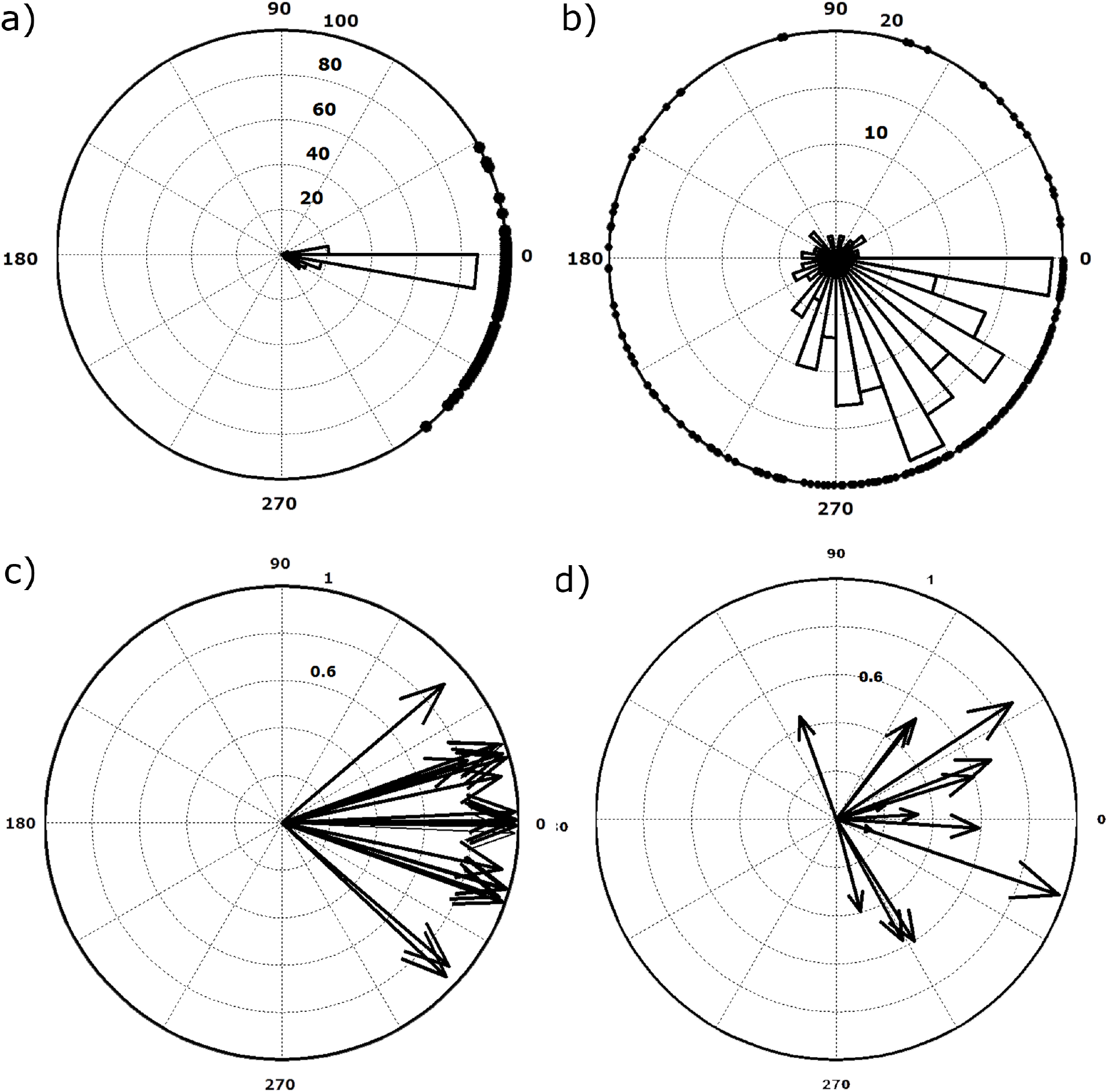
Synchrony in *Mecopoda* ‘Two Part Caller’. Top: Representative polar plots of timing relationships between a) trills and b) chirps of two calling males. Points on the circumference represent phase values. Histograms represent frequency of phase values in bins of 10°. Bottom: Mean phase vectors for c) trills and d) chirps across all interactions (N = 18 duets). Each vector represents the results from one male in a duet.

The verse period was significantly greater in duets than during solo calling for 9 out of 22 individuals (Fig. 4, Bonferroni corrected t tests, all Ps < 0.001, all ts < 6.02 and > −238.14) and significantly lower for 8 individuals (Fig. 4, Bonferroni corrected t tests, all Ps < 0.001, all ts < 16.91 and > −4.21). The mean absolute change in verse period from solos to duets was 276 (± 60.17 S.E.) ms. The mean difference in verse period between pairs of duetting males was 48.77(± 19.15 S.E.) ms. This was much less that the mean difference between the solo verse periods of each pair (317.77 ms ± 72.16 S.E) as might be expected because of the synchrony during duets.

**Fig. 4.**
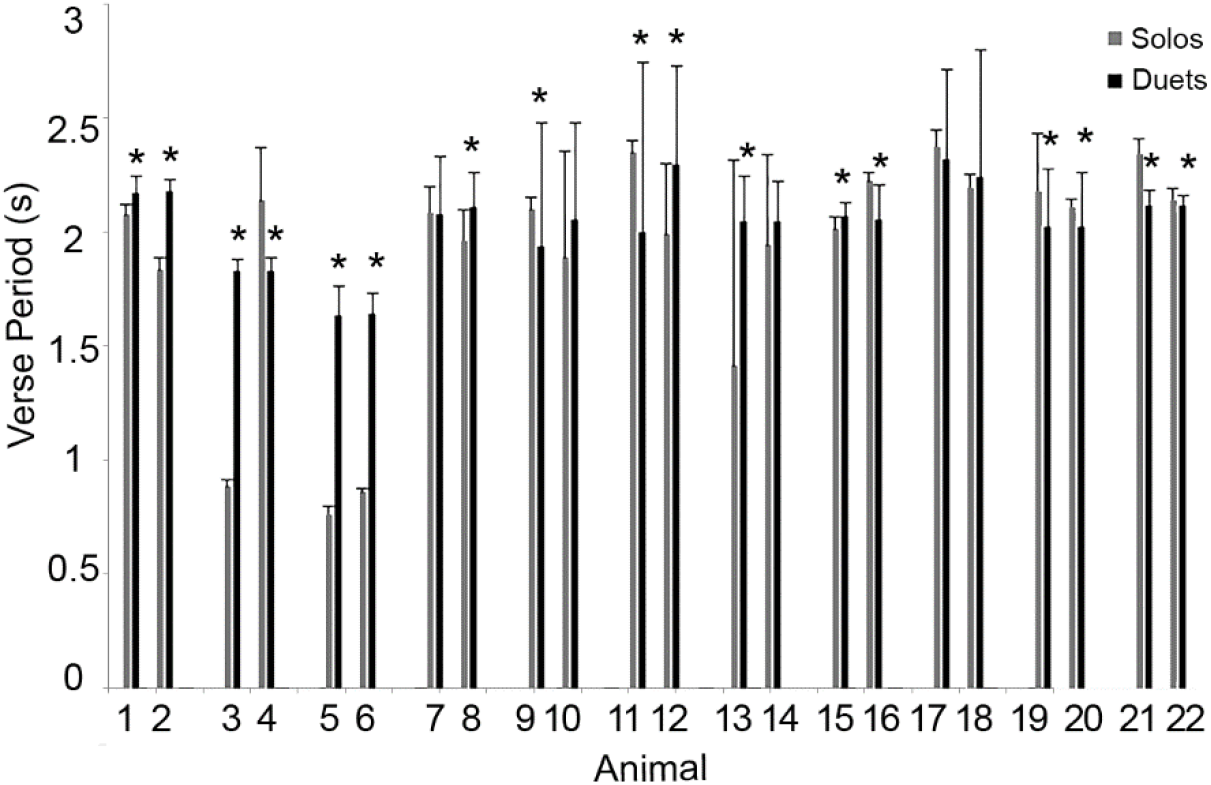
Change in verse period from solos to pairwise interactions. Grey bars represent values obtained during solos. Black bars represent values obtained during duets. Values obtained from pairs of males that interacted with each other are grouped together and the numbers below identify individual animals. Asterisks indicate significant differences between the values obtained in solos and the values obtained in duets (p < 0.0023).

### The effect of trill playback on trill Phase Response Curves

The phase response curves of individuals rose for about half the stimulus phase after which they had a slight dip below 1 before returning to a value of 1 (Fig. 5). This indicates a delay in response to early stimulus phases and an advance in response to the later ones. Both delay and advance were however, not pronounced in all except three individuals (Fig. 5, animals 2, 3 and 9). In a few individuals, there were some trills that were delayed by a large amount in response to trills in the middle stimulus phases (Fig. 5, animals 2, 4, 5 and 13). There was no effect of stimulus phase on the timing of subsequent trills.

**Fig. 5.**
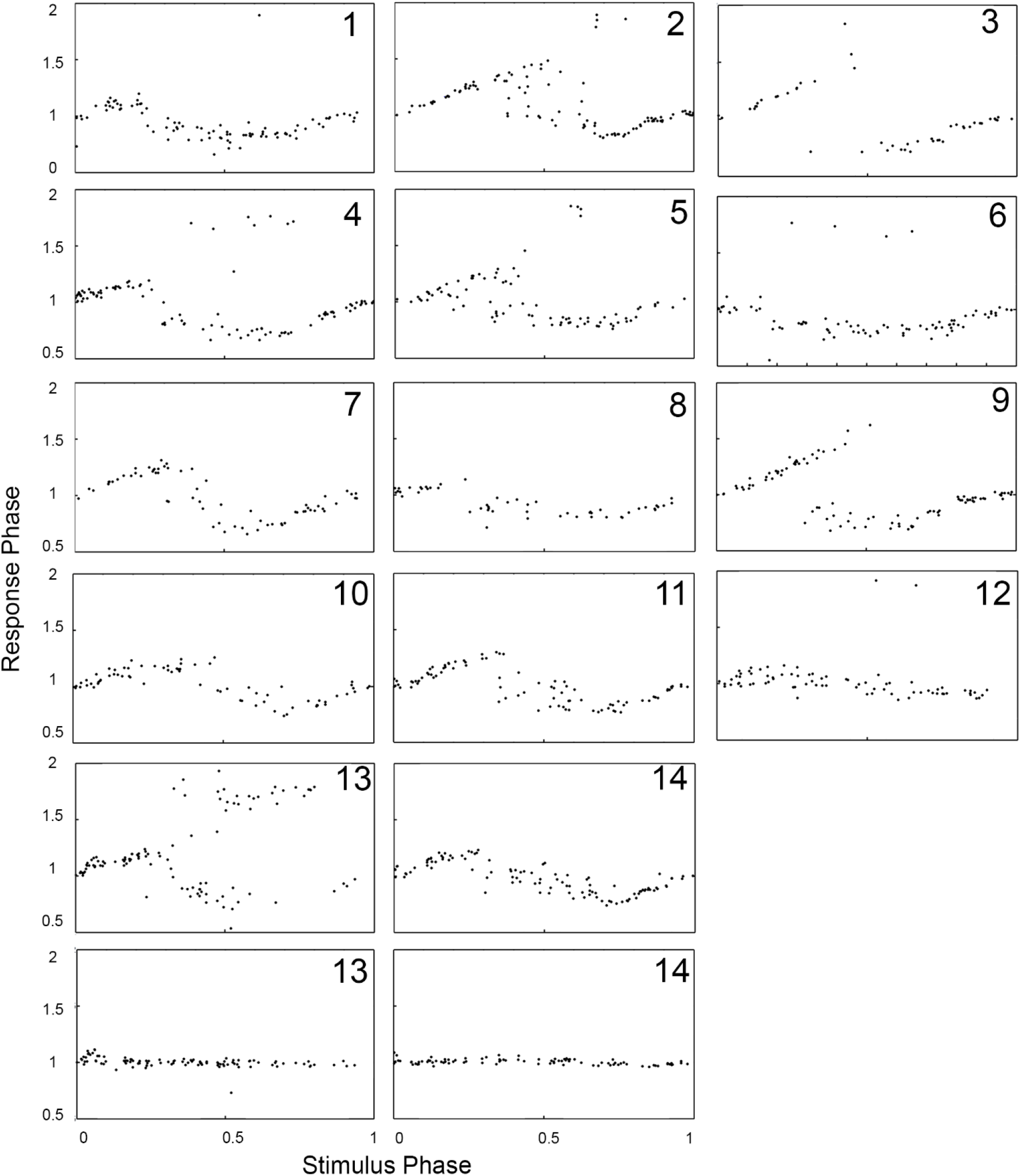
Trill Phase Response Curves obtained by plotting the response phase of trills in response to playback of stimulus trills. The last row depicts the phase response in the period following the disturbed period. Numbers refer to individual animals.

An analysis of verse periods before and after the playback of stimulus trills showed that in 6 out of 14 individuals, the average of three periods following the trill playback was significantly lower than the average of three periods preceding it (Bonferroni corrected t tests, all Ps < 0.003, all ts> 0.02). The mean difference between the periods before and after the stimulus trills was 64 (± 16 SE) ms. There was, however, no significant difference between the averages of periods preceding a stimulus trill and periods preceding the subsequent stimulus trill (t tests, all Ps > 0.68, all ts> −0.13 and < 0.42) with the mean differences between these two sets being 30 (± 17 SE) ms.

### The effect of trill playback on chirp Phase Response Curves

Since it was less likely that the onset of our playback trill would fall in the chirp section of the calling verse of the animal, we could not obtain individual Phase Response Curves for each animal. Instead we combined data across all animals to generate a single PRC. This PRC indicates that the playback of trills increases the period of the chirps if ending in the first half of the calling period (stimulus phase less than 0.5) (Fig. 6a). There was, however, no systematic increase of response phase with increasing stimulus phase which is often a characteristic of PRCs in other species.

**Fig 6.**
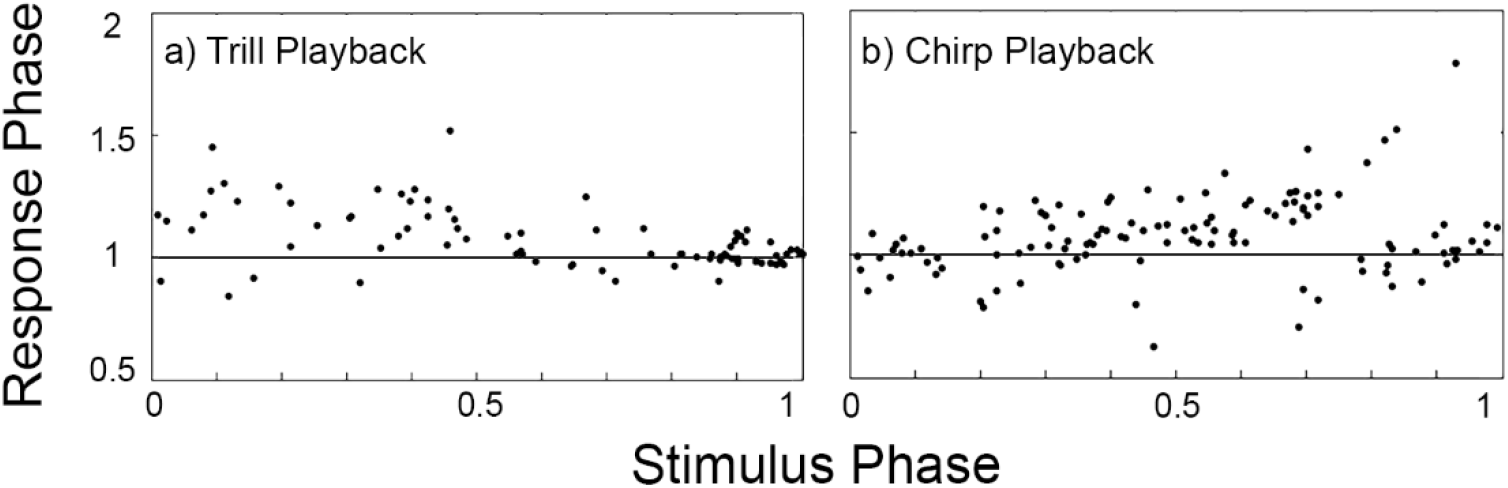
Chirp Phase Response Curves obtained by plotting the response phase of chirps in response to playback of a) stimulus trills and b) stimulus chirps. Both plots combine data from multiple animals.

One interesting aspect of trill playback, was that the number of chirps produced by the animal was significantly greater when the trill playback overlapped with the offset of the animal’s trill (paired t test, t = 4.83, P =3.27 *10^-4^, N = 14 males). The mean number of chirps produced was 2.74 (± 0.5 S.D.) when the playback overlapped the animal’s trill offset (N=14 males, 55.07 ± 15.37 playback trills per individual) and 2.10 (± 0.44 S.D.) when the playback did not overlap (N= 14 males, 30.36 ± 8.35 playback trills per individual).

### The effect of chirp playback on trill Phase Response Curves

Chirps played to individual males at random phases during their calls had no effect on the timing of the trills of the males (Fig. 7). The PRCs obtained were flat and the response phase stayed at a value of approximately 1 for all stimulus phases. The average of three verse periods before and after chirp stimuli did not significantly differ for 11 out of 12 individuals (t tests, all Ps > 0.12, all ts >−0.27 and <2.16) and the mean difference between periods before and after the stimuli was 32 (± 10 SE) ms.

**Fig. 7.**
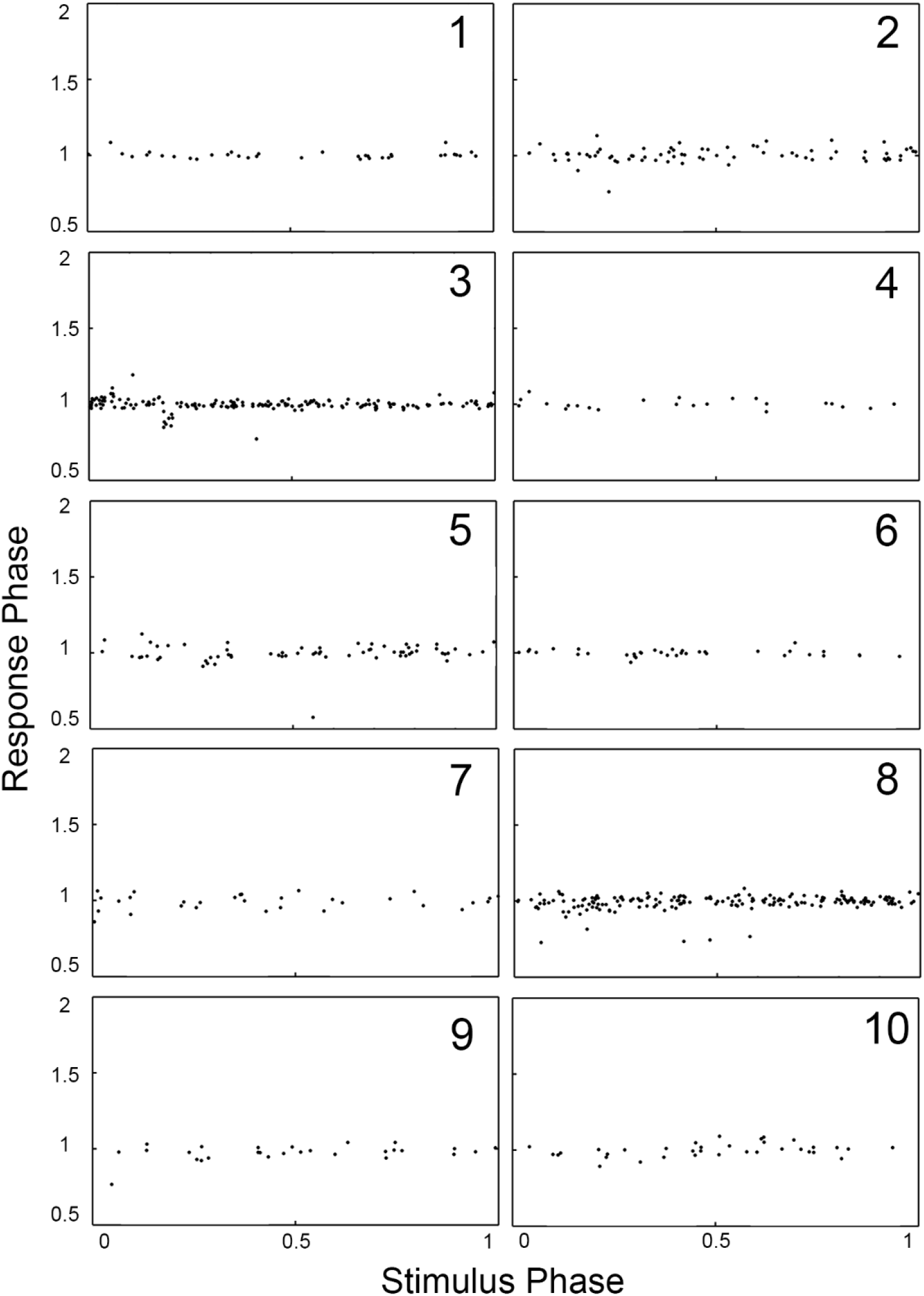
Trill Phase Response Curves obtained by plotting the response phase of trills in response to playback of stimulus chirps. The last row depicts the phase response in the period following the disturbed period. Numbers refer to individual animals.

### The effect of chirp playback on chirp Phase Response Curves

As discussed above, due to the lack of cases when the playback chirps overlapped the animal’s chirps, we were only able to obtain a combined PRC across all animals. This PRC described how the animals adjusted their chirp periods in response to playback of chirps (Fig. 6b). The response phase increased above 1 with increasing stimulus phase up to around a stimulus phase of 0.7. After a stimulus phase of around 0.7, the response phase was mostly below 1 and reach 1 at a stimulus phase of 1. Thus, the chirp periods were increased in response to chirps played back early in the chirp period, but were shortened in response to chirps played back later in the chirp period.

### Response to chirpless calls

11 out of 14 males synchronised their trills with those of the positive control stimulus played out to them (Fig. 8). The phase angles of the mean vectors obtained in these duets were different from a uniform distribution and were centred around 0° (V test, v = 9.36, P =2.0044*10^-4^). Three out of the 14 individuals, however, failed to synchronize their trills and instead had a phase angle closer to 90°.

**Fig. 8.**
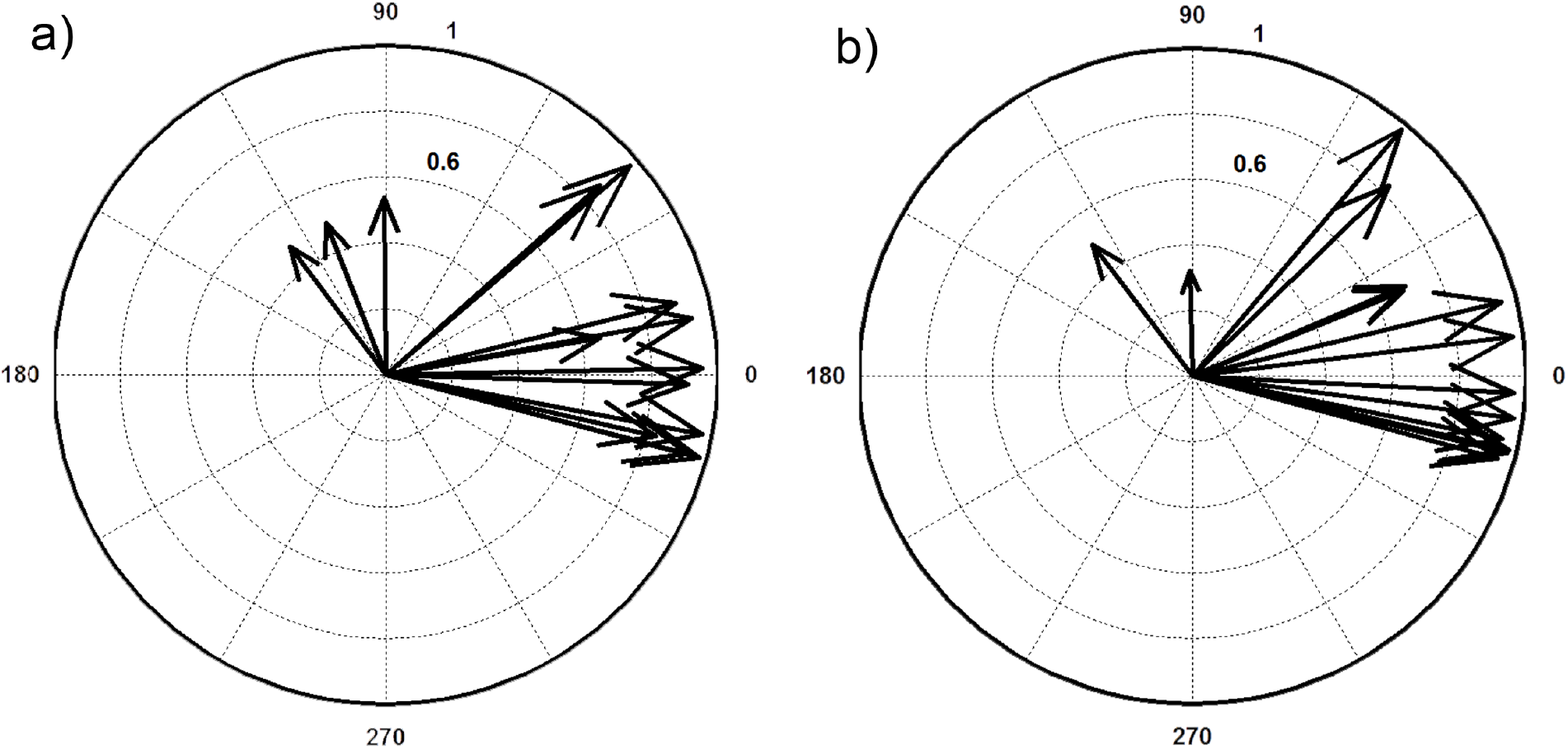
Mean phase vectors of trills during interactions with a) a positive control playback stimulus and b) a chirpless playback stimulus.

12 out of 14 males also synchronized their trills with those of the stimulus (chirpless calls). The phase angles of the mean vectors obtained in these duets were not uniformly distributed and were centred around 0° (V test, v = 10.42, P =4.11*10^-5^). Similar to the positive control duets, two out of 14 individuals failed to synchronize and had mean vector angles above 90°.

## Discussion

Our results show that individuals of the species *Mecopoda* ‘Two Part’, a katydid with a complex non-isochronous call, synchronize elements of their calls. The synchrony is governed by mechanisms similar to those in other synchronizing katydid species and gives us an insight into the oscillators underlying song production in this species.

### Phase Response Curve comparisons

Like several synchronizing katydid species, the Trill Phase Response Curve obtained for *Mecopoda* ‘Two Part Caller’ has both an ‘advance’ phase and a ‘delay phase’. Similar to *Mecopoda elongata,* both phases affect the immediate chirp and not the following chirp. This distinguishes the PRC from that of many other well studied species like the katydids *Neoconocephalus spiza* and the firefly *Pteroptyx cribellata* (Buck et al., 1981a; Greenfield and Roizen, 1993a). In these species, the immediate chirp is only delayed, and the advance phase only affects the subsequent chirp. Previous models such as the phase advance and phase delay models that have been used to explain the mechanisms of synchrony are thus insufficient to explain trill synchrony in this species. Unlike the PRCs predicted by these models, the trill PRC obtained for most individuals in *Mecopoda* ‘Two Part Caller’ is nonlinear and less pronounced. This PRC is perhaps more similar to that modelled for visual synchrony in other firefly species such as *Pteroptyx malaccae* (Ermentrout, 1991). Using one classification of firefly ‘pacemakers’ (Hanson, 1978), *Mecopoda* ‘Two Part Caller’ has a variable intrinsic chirp period with a small amplitude PRC. In this respect it differs from both previously studied species of the genus. Both *Mecopoda elongata* and *Mecopoda* ‘Chirper’ have high amplitude PRCs with two distinct arms and a break point around 70% into the period (Hartbauer et al., 2005; Nityananda and Balakrishnan, 2007). In fact, the chirp PRC (Fig 6b) we obtained for ‘Two Part Caller’ fits with this pattern and resembles the PRC previously obtained for *Mecopoda elongata*. In contrast, the trill PRC for most individuals consists of a continuous curve without any break points. Thus, the mechanism for adjusting chirp periods resembles that of other chirping species but the mechanism for adjusting trill periods is different.

Like ‘Chirper’, however, ‘Two Part Caller’ also has a variable intrinsic chirp period. In both species, the chirp period before external acoustic stimulation differs from that after, but the difference is lost if periodic stimulation is missing. This resembles a model for firefly synchrony (Ermentrout, 1991), which assumes an adaptable free-running period that enables synchrony. Oscillators that fit this model have a period that adapts to the period of the external stimulus until the two are in synchrony. This means that whether the period of the male in duets increases or decreases will also ultimately depend on the rate changes that occur in the duetting partner, as can be seen in our results from actual duets (Fig 4). If the period of the external stimulus or duetting partner is out of the range of adaptability, the model predicts phase-locking with a small phase difference instead of synchrony. This might be more like the results obtained in the positive control playback experiments of our study where all males lagged the stimulus by a specific amount and some had poor synchrony.

### A two-oscillator model of synchrony

In contrast to other species from which synchrony has been reported, *Mecopoda* ‘Two Part Caller’ has a call with more than one call component: a trill and a series of chirps. Models of oscillators in chirping species such as *Necoconocephalus spiza* assume a single oscillator that increases in level and is reset when an external chirp is heard (Greenfield and Roizen, 1993a). In ‘Two Part Caller’, each calling component could, however, differentially affect the production of calls. In separate playback experiments, we played back both trills and chirps to calling males and examined how they affected calling. The results of our experiments indicate that while trill playback affects the timing and synchrony of trills, chirp playback does not seem to affect them. This suggests that trill production is governed by an oscillator that is unaffected by the chirps. Playback of trills does affect the adjustment of chirp periods, as does playback of chirps. We therefore propose a dual-oscillator model in which the production of trills and chirps are governed by two different oscillators (Fig. 9). During solo calling, the trill oscillator suppresses the production of chirps for the duration of the trill, after which the chirp oscillator produces the chirps. While interacting with another calling male, the trills heard reset the trill oscillator as seen in the trill PRC. The chirps, however, do not reset the trill oscillator but affect the production of chirps by the chirp oscillator if heard in between trills, thus enabling the level of chirp synchrony seen. Such a model would also predict that when a male’s trills are suppressed by external trills, the suppression of the focal male’s chirps should be released, and we would expect more chirps to be produced in this case. Our results seem to bear this out, since more chirps are produced when external trills extend beyond a male’s trills, and thus presumably suppress further trill production.

**Fig. 9.**
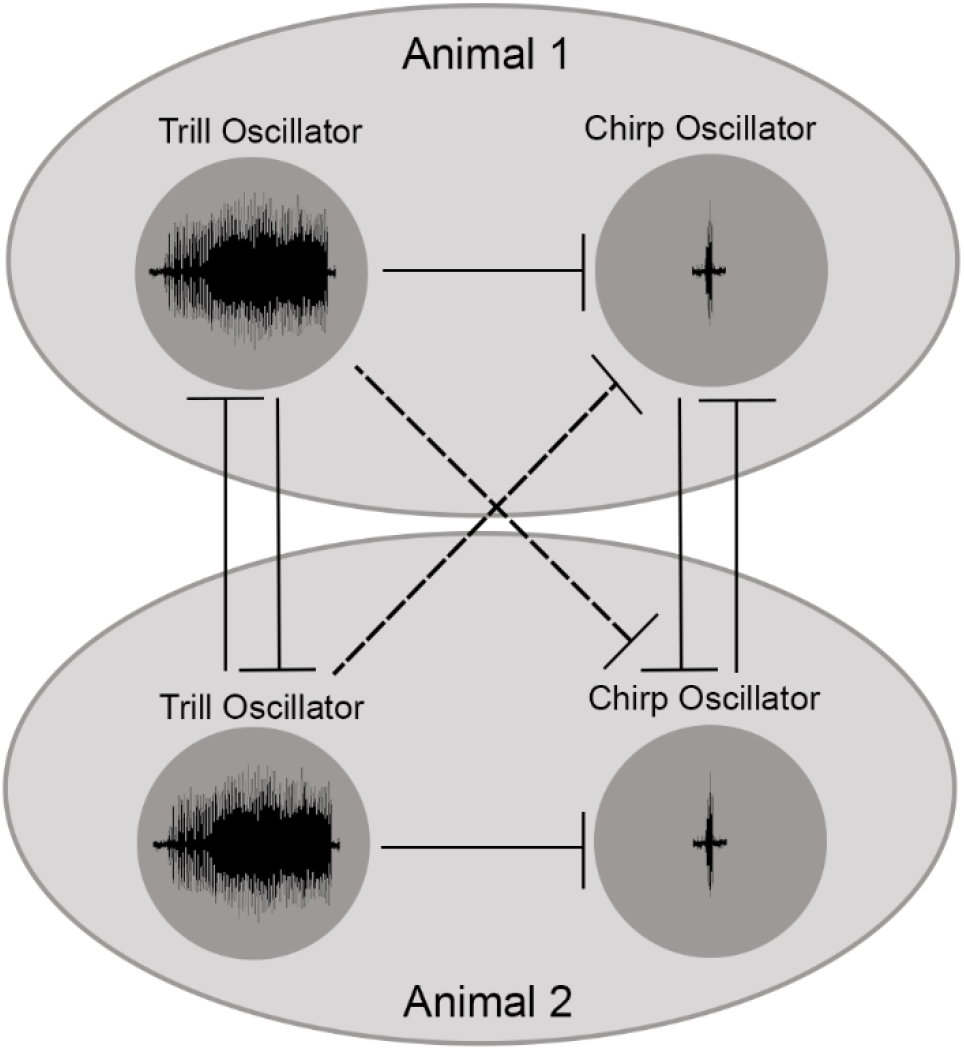
Two oscillator model for synchrony in *Mecopoda* ‘Two Part Caller’. Circles represent oscillators governing the production of trills and chirps. Larger ellipses represent each of two animals acoustically interacting with each other. Lines depict inhibitory processes. Inhibition between the trill oscillator and chirp oscillator of the other male is indicated as a dotted line to suggest a possibly different process.

### Advantages of leading trills and implications for the evolution of synchrony

In other synchronizing species of katydids, females have been shown to prefer chirps of males that lead the chirps of neighbouring males by a short window of time. In the species *Neoconocephalus spiza*, this preference could override other preferences for louder or longer chirps (Greenfield and Roizen, 1993a; Snedden and Greenfield, 1998). The mechanisms enabling chirp synchrony could have evolved as an evolutionarily stable strategy (ESS) in response to such a female preference (Greenfield and Roizen, 1993a; Greenfield et al., 1997). A female preference for leading chirps was also demonstrated in the species *Mecopoda elongata* (Fertschai et al., 2007). Such a preference may be due to the fact that leading chirps are preferentially represented in the auditory system of the female (Römer et al., 2002). The representation of the following chirp is inhibited for a specific time window following the leading chirp due to a process of contralateral inhibition. These results have important implications for the synchrony of trills in ‘Two Part Caller’. Given that longer calls with a higher duty cycle have greater inhibitory effects on the auditory system in katydids (Römer and Krusch, 2000), we would expect the long trills to have a large inhibitory effect on the auditory system of the female. If contralateral inhibition mechanisms similar to *Mecopoda elongata* operate in this species, a male with leading trills could therefore gain a huge advantage - the call of the following male would be inhibited for a long duration during which only the male with the leading trill would be heard. This could mean that females strongly prefer males with leading trills. Such a preference would create a strong selective pressure on the males to produce the same and could have driven the evolution of synchrony in this species in a process like that seen in *Necoconocephalus spiza*. An alternative evolutionary scenario could be that females prefer calls with chirps, and that the males need to synchronize the trills so that the chirps can be heard in the gaps. There might then be resemblances to the calls of the frog, *Physalaemus pustulosus*, which consists of a whine followed by ‘chucks’ where female prefer calls with chucks compared to calls without (Ryan, 1980).

To the best of our knowledge, this is the first report of acoustic synchrony of non-isochronous calls in an insect. This suggests that the ability to rhythmically respond to complex stimuli could be more widespread amongst animals than has previously been thought. However, it is important to clarify that the mechanisms underlying such abilities could be very different in different species. Our two-oscillator model suggests that the mechanisms of synchrony effectively reduce the differing rhythms in the call to two separate patterns of trills and chirps and the oscillators respond separately to each of these. This converts what might appear to be a complex pattern into a combination of two simpler mechanisms, which might not underlie rhythm perception in vertebrates, for example. Even amongst the closely related species of the katydid genus *Mecopoda*, it seems as if differing neural mechanisms could govern call synchrony. How these mechanisms have evolved and the selective pressures on them makes for a fascinating future area of research. Further studies, for example, are needed to shed light on female preferences for leading trills and the neural representation of these in the nervous system of ‘Two Part Caller’. It would also be important to investigate if, despite the similarities in synchronizing behaviour, the processes leading to the evolution of the behaviour in this species could differ, as has been suggested for the species *Mecopoda* ‘Chirper’ (Nityananda and Balakrishnan, 2009).

## Acknowledgements

VN is currently funded by a Biotechnology and Biological Sciences Research Council David Phillips Fellowship BB/S009760/1. We are grateful to the Ministry of Environment, Forests and Climate Change Government of India for funding this project. The equipment used in the study was funded by a DST-SERB grant to RB. We thank Dr Natasha Mhatre for help with writing the stimulus programs in Labview and Dr Hari Sridhar for help with fieldwork collecting the katydids. The experiments comply with the legal principles ofanimal care and animal welfare of the Government of India.

## Data Availability

All data supporting this paper can be accessed using the following link: https://doi.org/10.6084/m9.figshare.13293089.v1.

